# Effects of Oral Domoic Acid Exposure on Maternal Reproduction and Infant Birth Characteristics in a Preclinical Nonhuman Primate Model

**DOI:** 10.1101/440354

**Authors:** Thomas M. Burbacher, Kimberly S. Grant, Rebekah Petroff, Sara Shum, Brenda Crouthamel, Courtney Stanley, Noelle McKain, Jing Jing, Nina Isoherranen

## Abstract

Domoic Acid (DA) is a naturally-occurring excitotoxin, produced by marine algae, which can bioaccumulate in shellfish and finfish. The consumption of seafood contaminated with DA is associated with gastrointestinal illness that, in the case of high DA exposure, can evolve into a spectrum of responses ranging from agitation to hallucinations, memory loss, seizures and coma. Because algal blooms that produce DA are becoming more widespread and very little is known about the dangers of chronic, low-dose exposure, we initiated a preclinical study focused on the reproductive and developmental effects of DA in a nonhuman primate model. To this end, 32 adult female *Macaca fascicularis* monkeys were orally exposed to 0, 0.075 or 0.15 mg/kg/day DA on a daily basis, prior to and during pregnancy. Females were bred to non-exposed males and infants were evaluated at birth. Results from this study provided no evidence of changes in DA plasma concentrations with chronic exposure. DA exposure was not associated with reproductive toxicity or adverse changes in the physical characteristics of newborns. However, in an unanticipated finding, our clinical observations battery revealed the presence of subtle neurological effects in the form of intentional tremors in the exposed adult females. While females in both dose groups displayed increased tremoring, the effect was dose-dependent and observed at a higher frequency in females exposed to 0.15 mg/kg/day. These results demonstrate that chronic, low-level exposure to DA is associated with injury to the adult CNS and suggest that current regulatory guidelines designed to protect human health may not be adequate for high-frequency shellfish consumers.

**Highlights:** 1) Domoic acid acts as a tremoragen after chronic, low-dose oral exposure in adults.

2) Exposure across pregnancy does not result in maternal reproductive toxicity.

3) In-utero exposure does not adversely impact physical characteristics of exposed newborns.

4) Current regulatory guidelines may not adequately protect high-frequency shellfish consumers from DA-induced neurological injury.

## 1. Introduction

The six oceans of the world, the Arctic, Atlantic, Indian, Pacific and Southern, cover over 70% of the planet’s surface and comprise the largest ecosystem on earth. Within the ocean environment, there are thousands of microalgae species that play a positive role in producing thriving fisheries and rich feeding grounds for marine animals and seabirds. There are also a small number of microalgae that produce dangerous biotoxins and these species are referred to as Harmful Algal Blooms (HABs) (Centers for Disease Control and Prevention, 2017; Wells et al., 2015). HABs have been linked to large-scale killings of wildlife and are responsible for over 60,000 cases of human poisoning each year (Berdalet et al., 2016; Grattan et al., 2016b; Stommel and Watters, 2004). Domoic Acid (DA), the focus of this paper, is a naturally-occurring neurotoxin that is produced by harmful algae in the genus *Pseudo-nitzschia* (Bates et al., 1989; Garrison et al., 1992). A member of the kainoid class of excitatory neurotransmitters, DA is transferred up the food chain through bioaccumulation and elevated levels have been documented in mussels, oysters, razor clams, scallops, sardines, anchovies, crab and lobster. Shellfish and small finfish accumulate this toxin because they feed on the toxic algae that produce DA, and the primary route of exposure for humans is the consumption of contaminated shellfish meat and viscera (Washington State Department of Health). The algal blooms that produce this marine biotoxin are becoming more frequent, and in the summer of 2015, a *Pseudo-nitzschia* bloom off the Western coast of the United States was so massive it extended from the central California coast to the Alaska Peninsula and triggered an unprecedented number of beach closures to shellfish harvesting (McCabe et al., 2016). Warm ocean conditions and nutrient enrichment from seasonal upwelling have been identified as factors that are driving the increased occurrence and severity of these damaging blooms (McKibben et al., 2017; Zhu et al., 2017).

Exposure to DA can cause illness and death in humans and animals and is a growing public health concern in many coastal dwelling communities (Lefebvre and Robertson, 2010; Ramsdell and Gulland, 2014). The largest human episode of DA poisoning occurred on Prince Edward Island in Canada with 153 cases of acute intoxication and 4 deaths after patients consumed a meal of contaminated mussels (Perl et al., 1990a, 1990b). The outbreak in Canada provided vital data on acute, high-level exposure, and estimated patient exposures ranged from 60 to 290 mg DA/patient or about 1 to 5 mg/kg for a 60 kg person (Jeffery et al., 2004; Perl et al., 1990b). Subjects with overt symptoms of DA poisoning exhibited effects characterized by initial gastrointestinal illness (e.g. nausea, vomiting, diarrhea) that in some cases, evolved into a spectrum of responses ranging from agitation/confusion to hallucinations, memory loss, seizures and coma (Teitelbaum et al., 1990b, 1990a). The term *Amnesic Shellfish Poisoning* originated from clinical observations that memory functions were significantly impaired in some DA-sickened patients, even after recovery from overt signs of illness. Patient autopsies revealed neuronal loss and necrosis that was concentrated primarily in the amygdala and hippocampal regions of the brain. While there have been no significant, verified episodes of human DA poisoning since 1987, rising DA levels in ocean waters have been associated with the large-scale killing of marine mammals and seabirds (Lefebvre et al., 2016, 2002). It has been suggested that ocean-dwelling wildlife are acting as sentinel species for the human health risks associated with exposure to this dangerous and increasingly widespread neurotoxin (Backer and Miller, 2016; Bossart, 2011). Because over 123 million Americans live in a county directly on the shoreline (US Department of Commerce and National Oceanic and Atmospheric Administration), the physical well-being of marine mammals who inhabit the same coastal environments as humans is vitally important to understanding the biological threat that DA poses to human health.

In an effort to protect human health from the adverse effects of acute high-level DA exposure,, regulatory and environmental protection agencies have established a threshold of 20 ppm in razor clam meat as the level at which coastal beaches are closed to commercial and recreational shellfish harvesting (Mariën, 1996; Wekell et al., 2004). This level is based on estimates of DA exposure and effects from the Prince Edward Island poisoning. Researchers and environmental health agencies have also proposed various estimates of DA intake that protect public health. Proposed estimates for repeated exposure to DA or the Tolerable Daily Intake (TDI) vary from 0.075 to 0.1 mg/kg/day. These values are based on data from the Prince Edward Island poisoning or results from laboratory studies using nonhuman primates (FAO, 2004; Toyofuku, 2006; Marien, 1996). On the Western coast of the United States, more frequent algal blooms are resulting in the persistent contamination of shellfish with detectable levels of DA that do not reach the 20 ppm threshold. Individuals who regularly harvest/ consume shellfish with elevated but permissible levels of DA (under 20 ppm) are being chronically exposed to this marine biotoxin (Ferriss et al., 2017). Coastal-dwelling tribal nations are at particularly high-risk for long-term, low-dose exposure because of their reliance on the ocean as an essential economic resource, foundation of cultural identity and year-round food source. A dietary survey of tribal members living on three Native American reservations in Washington State found that 84% of respondents indicated recent razor clam consumption (Boushey et al., 2016; Fialkowski et al., 2010). The impact of chronic DA exposure was recently studied in these communities and exposure (as measured by the consumption of razor clams) was associated with decrements in memory that were serious enough to affect everyday living skills (Grattan et al., 2018, 2016a). This striking research finding suggests that chronic exposure to low-level DA from harvested shellfish is not without risk to the adult central nervous system. The growing significance of DA as an environmental threat demands that contemporary exposure scenarios be considered in the development of regulatory thresholds to protect human health. Very little is known about the impact of chronic, lower-level oral exposure and this is particularly the case for sensitive subgroups such as the developing fetus, infants and elders.

While preclinical work has been extremely valuable in defining the health risks of DA exposure, most studies have focused on the effects of acute, high-dose exposures in adult subjects using non-oral routes of exposure such as intraperitoneal (IP) and intravenous (IV) treatment (Grant et al., 2010). Reported signs of DA toxicity in mature monkeys and rodents are similar to those observed in the human poisoning cases and effects include hippocampal damage, adverse neurological effects (e.g. seizures) and the disruption of cognitive processes (Lefebvre et al., 2017; Scallet et al., 2004, 1993, Tryphonas et al., 1990a, 1990b). The reproductive and developmental effects of DA exposure have not been studied in humans or nonhuman primates, but the preclinical rodent literature suggests a marked fetal vulnerability (Dakshinamurti et al., 1993; Xi et al., 1997). Consequences of early life exposure include hippocampal damage (sometimes progressive in nature), seizure disorders and persistent changes in a number of behavioral domains, particularly memory, social behavior and emotional reactivity (see Doucette and Tasker, 2016 for review). Adverse effects in young animals are observed at levels of exposure up to 40 times lower than those required to produce neurotoxicity in adults.

The protection of human health, particularly in vulnerable groups such as developing fetuses, will depend upon our understanding of this chemically-complex compound when ingested orally at low doses on a chronic basis. To address this growing public health concern, we initiated a longitudinal, birth-cohort study to investigate the maternal reproductive and offspring developmental effects of chronic, low-level, oral DA exposure in nonhuman primates. Preclinical model systems offer the ability to control many of the confounding environmental variables that can adversely impact sensitive outcome measures in human epidemiological studies and can provide an important connection between brain and behavior (Thompson et al., 2009). In this investigation, we selected the macaque monkey to model human exposure effects over pregnancy because of shared reproductive attributes between the two species (e.g. similarities in the morphology and physiology of the ovaries, the presence of discoid, hemochorial placentas and prolonged pregnancies (Buse et al., 2003; Carter, 2007; King, 1993). To increase the translational impact of study results, DA doses were carefully selected to mimic chronic exposure at or near the proposed Tolerable Daily Intake (TDI) for humans (Costa et al., 2010; Mariën, 1996; Toyofuku, 2006). This manuscript presents the first data from primates that addresses the maternal and fetal risks of this biological marine contaminant and provides insight into the adequacy of current environmental regulations to protect the human health in high-frequency shellfish consumers.

## 2.0 Materials and Methods

### 2.1 Chemicals and Reagents

DA was purchased from BioVectra (Charlottetown, PE, Canada). Certified calibration solution for DA was purchased from National Research Council Canada (Ottawa, ON, Canada). Optima grade water, methanol, acetonitrile, and formic acid used for bioanalytical assays were purchased from Fisher Scientific (Pittsburgh, PA).

### 2.2 Subjects

Thirty-two adult female and 3 adult male *Macaca fascicularis* were acquired for the study. The females were pseudo-randomly assigned to either a control group (N=10), a 0.075 mg/kg/day DA group (N=11) or a 0.15 mg/kg/day DA group (N=11) to establish groups with similar ages and weights. Females ranged in age from 5.5 to 11 years with an average age of approximately 7 years across the groups. Females ranged in weight from 2.8 to 4.2 kg with an average weight at assignment of approximately 3.5 kg across the groups. All females were nulliparous.

### 2.3 Study Design

All animal procedures strictly followed the Animal Welfare Act and the Guide for Care and Use of Laboratory Animals of the National Research Council. All protocols were approved by the University of Washington Institutional Animal Care and Use Committee to meet the highest standards of ethical conduct and compassionate use of animals in research. During the study, adult females were housed in individual communicating cages that were equipped with grooming bars that allowed adjacent females contact 24 hours/day. Frozen treats, fresh fruit and vegetables, music, movies and toys were provided on a routine basis. The adult males were housed individually in a separate room away from the females and provided the same food and enrichments. Animal housing rooms was kept at ~75 degrees and lights were on a 12 hour on and 12 hour off schedule.

The study design was based on previous reproductive and developmental toxicology studies conducted in our laboratory (Burbacher et al., 2004, 1987). The study included a training period where protocols were implemented to train females to drink from a syringe, collect blood for DA analysis without sedation, evaluate general health status (weights, medications), observe signs of DA toxicity (Clinical Observations), and detect menstrual bleeding. Once females were trained, experimental protocols were conducted for 2 months without DA exposure (Baseline Period). During this period females were orally dosed on a daily basis with the dosing vehicle solution alone (5% sucrose in water). Daily, oral exposure to DA was then initiated for the 2 DA exposure groups (the controls continued to receive the dosing vehicle alone) and dosing continued for another 2 months prior to breeding and then throughout breeding (Pre-Pregnancy DA Exposure Period) and pregnancy (Pregnancy Period).

Experimental protocols continued throughout these periods. Males were not exposed to DA.

### 2.4 Daily DA Dosing Protocol

Dosing solutions in 5% sucrose were prepared fresh every week. Briefly, sucrose was weighed to prepare a 5% (w/v) sucrose solution in tap water. DA powder was weighed and dissolved in the 5% sucrose solution to produce a 1 mg/mL DA solution. The DA solution was sonicated for 15 minutes to ensure complete dissolution and filtered subsequently using a syringe filter (0.2 microm, 25mm). The filtered 1 mg/mL DA solutions were subsequently diluted according to the animals’ weights with filtered sucrose solution. The prepared individual dosing solutions were stored in 16 mL glass vials at 4°C until use. Prior to pregnancy, females were dosed according to their last weekly weight. During pregnancy, females were dosed according to their last pre-pregnant weight. Syringes were labeled with only the animal number and week to ensure all testers remained blind to the exposure groups. Dosing occurred at approximately the same time every morning (~9 AM), seven days a week. Dosing solutions were analyzed for DA content by LC-MS/MS to confirm DA concentrations (Shum et al., 2018). Females were fed Lab Diet High Protein Monkey Diet biscuits approximately 2 hours before the daily DA dosing and again approximately 5 hours after dosing.

### 2.5 Blood Collection and DA Analysis Protocols

Blood samples were collected every two weeks for DA quantification across the study. Blood collections occurred approximately 5 hours after dosing with sucrose during the baseline period and alternated between collections at 5 hours post DA dosing or 24 hours post DA dosing during the exposure period. Blood samples were also collected at delivery. Sedation was not used for the bi-weekly blood collection because the animals were trained to provide their leg for the blood draw. Blood was drawn from the great saphenous vein into sodium heparin tubes and centrifuged at 3,000 x*g* for 15 minutes to isolate plasma for analysis. Plasma was stored at −20°C until analysis. DA was measured using a recently developed and validated LC-MS/MS method (Shum et al., 2018). The lower limit of detection (LOD) was 0.16 ng/mL and the lower limit of quantitation (LLOQ) was 0.31 ng/mL of DA in monkey plasma.

### 2.6 Maternal Health Assessments

#### Maternal Weight

All females were weighed once per week during the investigation. Animals were trained to voluntarily enter a transfer box which was placed on a digital scale where values (minus the weight of the transfer box) were recorded in kilograms.

#### Maternal Medications

Females were observed for signs of diarrhea, lethargy, weight loss, on a daily basis by study personnel. Females exhibiting signs of illness were reported to the attending veterinarian and medications were prescribed if necessary. Information regarding the type, dosage, and duration of various medications were recorded.

#### Clinical Observations

A Clinical Observation Test developed previously (see Burbacher et al., 1999, 2004) was used to provide a rapid evaluation of the visual functioning and motor coordination of the females. Females were typically observed at least 3 times per week, ~5 hours after dosing. Testers initially presented a small piece of fruit approximately 6 to 10 inches from the cage and recorded whether or not the females visually oriented to the fruit. The fruit was then moved to the left or right and recorded whether or not the females visually followed the fruit. Testers recorded the absence or presence of visual orientation as well as whether a full, partial or no follow was observed during the trial. Testers then placed the small piece of fruit or cereal in the palm of their hand and held it approximately 6 to 8 inches from the cage so the females had to reach out and pick up the treat. The tester observed whether or not hand and/or arm tremors were observed during the reach. Fine motor coordination was also evaluated by noting whether or not the female was able to retrieve the small treat. Three “reaching” trials were presented during a session. In addition to the above, testers recorded whether they observed vomiting, scratching, lethargy or any other abnormal behaviors during the observation test. All testers were trained to be reliable on this procedure (reliability criteria = 80%) and testers were blind to the exposure history of the females. Testers and care staff also recorded any observations of vomiting, diarrhea or any abnormal behaviors throughout the day and reported these observations to the appropriate clinician.

### 2.7 Maternal Reproductive Assessments

#### Tracking Menstrual Bleeding

Using positive reinforcement and behavioral shaping, animals were trained to voluntarily present their perineum for visual inspection to detect menstrual bleeding. A flashlight was used to help examiners and serve as a cue and animals were rewarded with a treat after turning, lifting their tail and allowing a brief visual inspection. The first day that bleeding was detected was consider Day 1 of the menstrual cycle. Observations for menstrual bleeding occurred seven days a week.

#### Breeding

Females were bred with one of three nonexposed males after 4 menstrual cycles were observed (2 prior to DA exposure and 2 after DA exposure had begun). Females were bred up to 4 hours and were checked for the presence of semen (providing evidence that breeding occurred) after each breeding. For the initial 3 breedings, females were placed with 1 of the males for approximately 9 days during mid-cycle. For breedings after that, females were housed with a male during the entire cycle. A maximum of 7 breedings occurred. Pregnancy confirmation was determined with a progesterone blood assay or ultrasound.

#### Monitoring Pregnancy and Delivery

As mentioned above, females were observed on a daily basis by study personnel. Beginning at approximately 135 days of pregnancy, females were observed for the onset of labor during the evening (6 pm to 6 am) every 30 minutes via infrared camera by nursery staff. When signs of active labor were observed, the delivery team was summoned to the laboratory in order to separate the newborn at birth to begin our postnatal assessments. Adult females were sedated with ketamine (IM, 5 mg/kg) after they delivered and the infant was placed in a preheated incubator for transport to a specialized nursery for evaluation. Adult females were monitored by nursery staff on camera until fully recovered from sedation. If delivery complications occurred (e.g. prolonged labor, abnormal delivery position), the veterinary staff was immediately notified that assistance was needed. Veterinarians with extensive primate experience made all clinical decisions related to labor and delivery and were fully empowered to determine when the labor process was not progressing normally and a cesarean section was required to protect the well-being of the infant.

### 2.8 Newborn Procedures

All examiners were blind to the prenatal exposure history of the infants and had been extensively trained on all data collection measures.

#### Newborn Weights and Anthropometrics

Newborns were weighed and an anthropometric exam was administered to measure crown-rump length and head circumference, width and length according to procedures previously described (Burbacher et al., 2004). Briefly, crown-rump length was measured by placing the infant in a supine position on a metal sliding scale with its rump against the base of the scale and its crown against a sliding barrier. The distance between the top of the infant’s head and the rump was measured. Head circumference was measured with use of a measuring tape that was placed around the infant’s head below the brow ridge, along the nuchal crest and above the ears, thus covering the occiput. Head width and length were measured using spreading calipers. Head width was measured using landmarks just above and in front of the ears, which provided a measure of the two widest points of the skull. Head length was measured using the length from the area between the eyes to the occiput. Two measurements (in millimeters) were made by reliable testers and the mean measurement was used for analysis.

#### Newborn Health Status (Apgar) and Screening for Congenital Defects

A procedure to measure newborn health status modeled after the human Apgar exam (Apgar, 1953) was administered twice, once within 30 minutes after birth and then again ~ 1 hour after birth. The exam includes measures of heart rate, respiration, temperature, muscle tonicity, physical activity and skin color. A screening exam for physical birth anomalies was also undertaken within the first 24 hours that focused on the skin (pigmentation, rash, bruises, edema), head (molding, fontanel, depressions, overriding suture), face (symmetry, mouth, eyes, ears, teeth), trunk (sternal retraction, spine, breasts, neck), genitals (vagina, anus, penis, scrotum) and extremities (digits, limbs).

#### Blood Collection

Blood samples were collected from the femoral vein of offspring on postnatal day 1. Blood was processed according to the procedures described above.

## 3.0 Statistics and Results

### *3.1* Maternal DA Exposure

The duration of DA exposure for the adult females ranged from 79 to 233 days at conception and 232 to 403 days at delivery and was largely dependent on the number of breedings required for a conception (see Table 1). While the overall range is quite large, the results of one-way analysis of variance (ANOVA) tests did not indicate a significant difference in the number of days exposed to DA (or sugar solution) at conception and delivery as a function of DA exposure group (conception, F(2,26) = 0.67, p=0.52; delivery, F(2,26) = 0.55, p=0.58).

**Table 1:** **Mean ± SE (Range) # of Days Exposed to DA at Conception^1^ and Delivery**

| DA Exposure Group | Conception | Delivery |
| --- | --- | --- |
| Controls (N=10) | 113±13 (82-194) | 274±14 (232-361) |
| 0.075 mg/kg/day (N=10) | 132±19 (84-233) | 294±19 (239-403) |
| 0.15 mg/kg/day (N=11) | 139±17 (79-232) | 298±18 (233-387) |

<sup>1</sup> End of breeding for females who did not conceive
Note: No significant DA exposure group differences in # of days exposed to DA at conception or delivery

### Maternal Plasma DA Concentrations

The results of the bi-weekly blood draws are shown in Figure 1 and Table 2. Samples that were above the LOD (0.16 ng/mL) but below the LLOQ (0.31 ng/mL) for DA were designated as Below Limit of Quantitation (BLQ) and assigned the value 0.24 mg/ml (half way in between the LOD and the LLOQ). DA was not detected in any plasma samples during the baseline period. Once DA exposure began, plasma DA concentrations varied with continuous DA exposure prior to pregnancy. Plasma DA concentrations at 5 hours post-dosing ranged from 0.24 ng/ml to 5.26 ng/ml for the 0.075 mg/kg/day exposure group and 0.37 ng/ml to 10.01 ng/ml for the 0.15 mg/kg/day exposure group. Plasma DA concentrations at 24 hours post-dosing ranged from 0.24 ng/ml to 2.36 ng/ml for the 0.075 mg/kg/day exposure group and 0.24 ng/ml to 5.61 ng/ml for the 0.15 mg/kg/day exposure group (see Figure 1). The results of one-way ANOVA tests using the mean individual blood DA concentrations for each female prior to pregnancy (or at the end of breeding for females who did not conceive) indicated that plasma DA concentrations were significantly higher for the 0.15 mg/kg/day exposure group compared to the 0.075 mg/kg/day exposure group at both sample time-points (5 hour sample, F(1,20) = 28.48, p<0.01; 24 hour sample, F(1,20) = 27.66, p<0.01).

Plasma DA concentrations continued to vary during pregnancy. Plasma DA concentrations at 5 hours post-dosing ranged from 0.24 ng/ml to 5.26 ng/ml for the 0.075 mg/kg/day exposure group and 0.37 ng/ml to 9.16 ng/ml for the 0.15 mg/kg/day exposure group. Plasma DA concentrations at 24 hours post-dosing ranged from 0.24 ng/ml to 2.36 ng/ml for the 0.075 mg/kg/day exposure group and 0.24 ng/ml to 4.63 ng/ml for the 0.15 mg/kg/day exposure group during pregnancy (see Figure 1 and Table 2). The results of one-way ANOVA tests using the mean individual blood DA concentrations for each female during pregnancy again indicated that plasma DA concentrations were significantly higher for the 0.15 mg/kg/day exposure group compared to the 0.075 mg/kg/day exposure group at both sample time-points (5 hour sample, F(1,18) = 18.27, p<0.01; 24 hour sample, F(1,18) = 24.14, p<0.01). Finally, a repeated measures ANOVA comparing the mean DA plasma concentrations prior to pregnancy with those during pregnancy indicated a significant difference for the 5- hour sample F(1,18) = 5.53, p=0.03). Five-hour DA blood concentrations were significantly lower during pregnancy for both DA exposure groups. No significant differences were observed for the 24-hour sample.

**Table 2:** **Mean ± SE (Range) Maternal Plasma DA Concentrations (ng/ml) Before and During Pregnancy**^1^

| DA Exposure Group | Before Pregnancy<sup>2</sup> |  | During Pregnancy |  |
| --- | --- | --- | --- | --- |
|  | 5-Hour Sample | 24-Hour Sample | 5-Hour Sample | 24-Hour Sample |
| 0.075 mg/kg/day (N=10) | 1.31±0.18<br>(0.62-2.76) | 0.49±0.07<br>(0.24-0.98) | 0.93±0.22<sup>3</sup><br>(0.46-2.57) | 0.50±0.07<sup>3</sup><br>(0.28-1.04) |
| 0.15 mg/kg/day (N=11) | 3.42±0.35<br>(1.48-5.35) | 1.65±0.21<br>(0.61-2.47) | 2.93±0.38<br>(0.69-4.69) | 1.50±0.17<br>(0.55-2.39) |

<sup>1</sup> Mean of individual mean values for each female. For 0.075 mg/kg/day DA exposure group, mean calculations include 69 BLQ samples (35 24-hour samples before pregnancy, 29 24-hour samples during pregnancy and 5 5-hour samples during pregnancy). For 0.15 mg/kg/day DA exposure group, mean calculations include 9 BLQ samples (3 -24-hour samples before pregnancy and 6 24-hours samples during pregnancy)
<sup>2</sup> End of last breeding for females who did not conceive
<sup>3</sup> N=9 during pregnancy
Note: Maternal DA plasma concentrations significantly higher in 0.15 mg/kg/day DA exposure group prior to and during pregnancy ( $p < 0.01$ , all tests). 5-Hour Maternal DA plasma concentrations significantly lower during pregnancy ( $p = 0.03$ ).

**Figure 1.**
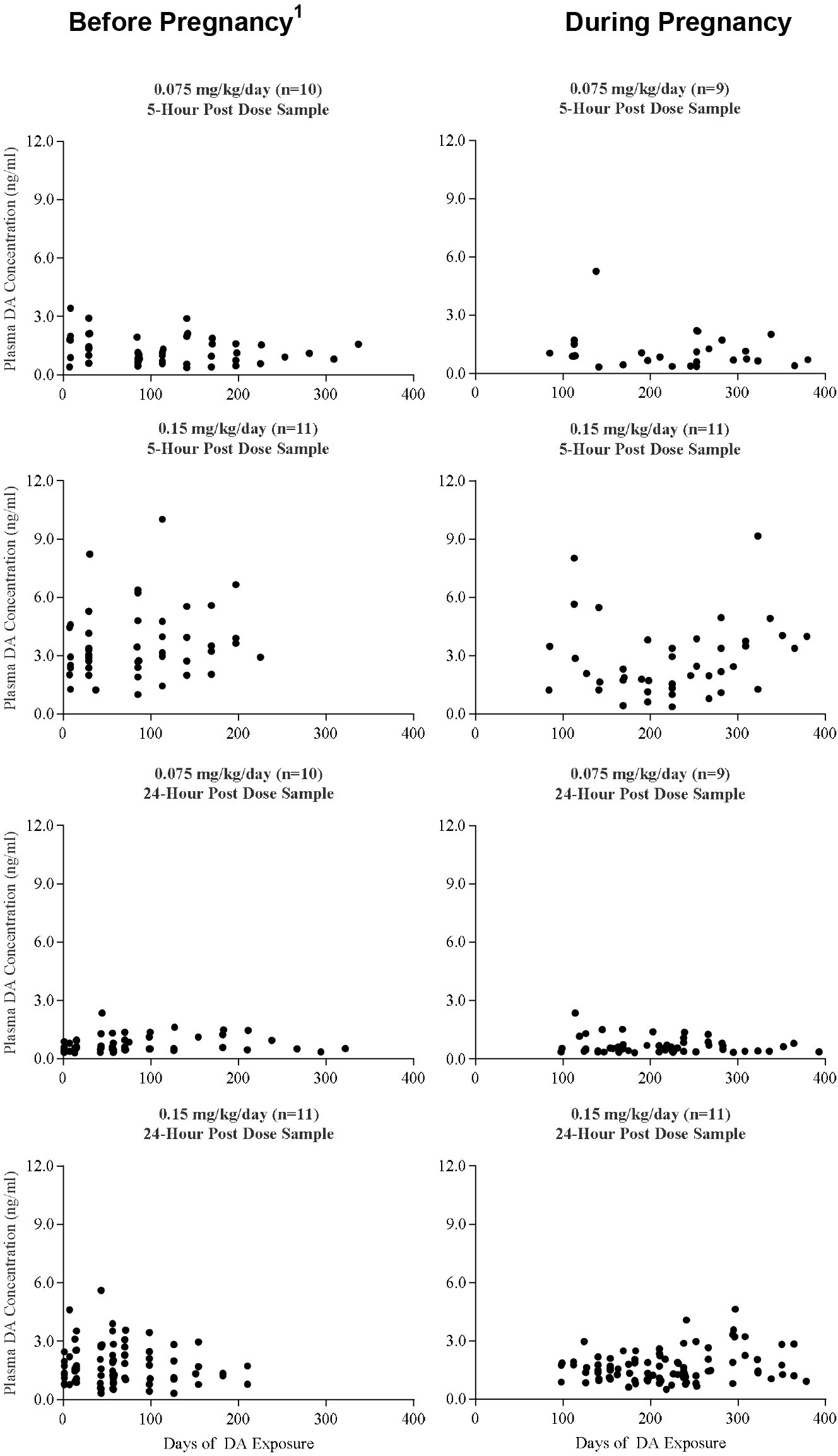
**Maternal plasma DA Concentration (ng/ml) Before and During Pregnancy**

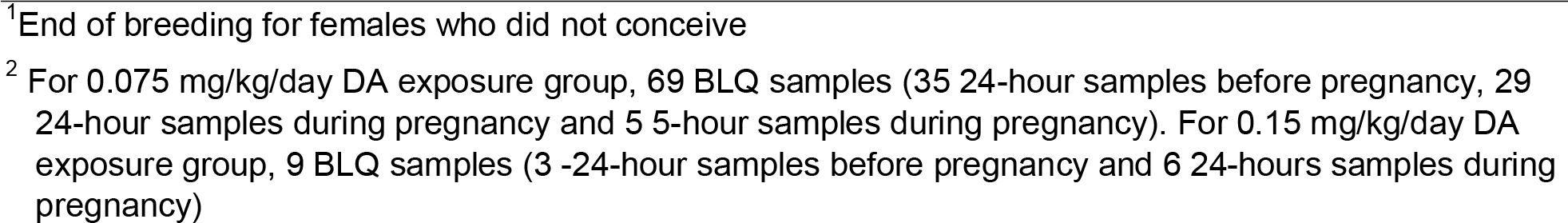

## 3.2 Maternal Health Assessments

### Maternal Weight

Maternal weights were similar across the DA exposure groups throughout the study (see Table 3). Except for 1 female, weights changed little once DA exposure began. One female in the 0.075 mg/Kg/day DA exposure group was removed from the study during the breeding period after she exhibited weight loss and amenorrhea. The results of a two-way ANOVA comparing the weights of the 3 DA exposure groups across the baseline and pre-pregnancy DA exposure periods did not indicate a significant difference due to DA exposure (F(2,28) = 0.46, p=0.67), or an interaction of DA exposure and exposure period (F(2,28) = 0.79, p=0.46). A main effect of exposure period was observed, indicating an increase in the weight of females during the pre-pregnancy period (F(1,28) = 18.83, p<0.01). For those females who conceived, the results of a one-way ANOVA did not indicate a significant difference in weight during pregnancy due to DA exposure (F(2,26) = 0.78, p=0.47). Weight gain during pregnancy was also similar across the 3 DA exposure groups (F(2,26) = 0.63, p=0.54).

**Table 3:** **Mean ± SE (Range) Weights (Kg) of Females Prior to and During DA Exposure and Weight Gain (Kg) during Pregnancy**

| DA Exposure Group | No DA Exposure | DA Exposure |  |
| --- | --- | --- | --- |
|  | Baseline Weights<sup>1</sup> | Pre-Pregnancy Weights<sup>1,2</sup> | Pregnancy Weight Gain |
| Controls (N=10) | 3.50±0.10 (2.98-3.91) | 3.63±0.13 (3.08-4.23) | 1.72±0.17 (1.16-2.88)<sup>3</sup> |
| 0.075 mg/kg/day (N=10) | 3.58±0.18 (2.89-4.77) | 3.85±0.22 (3.04-5.05) | 1.95±0.21 (1.13-2.97) |
| 0.15 mg/kg/day (N=11) | 3.64±0.13 (3.05-4.29) | 3.86±0.18 (3.19-4.81) | 1.62±0.24 (0.35-2.56)<sup>3</sup> |

<sup>1</sup> 4 month baseline period, ~3 to 8 month Pre-Pregnancy period
<sup>2</sup> End of last breeding for females who did not conceive
<sup>3</sup> N=9
Note: No significant DA exposure group differences in weights of females during baseline or pre-pregnancy DA exposure periods. No significant DA exposure group differences in weight gain of females during pregnancy.

### Maternal Medications

All females were sedated with approximately 10 mg/kg IM ketamine for mandatory tuberculosis testing approximately 1 month before breeding. During the early stages of the study (prior to pregnancy), six females required treatment for minor injuries incurred during socialization. These treatments generally included at least one pain medication (ketoprofen, meloxicam, buprenorphine or lidocaine) and at least one antibiotic (cephalexin, cefazolin, or penicillin). In addition, two females required treatment during pregnancy. One was treated with acetaminophen and a single dose of penicillin and buprenorphine. The other female was treated with acetaminophen, buprenorphine and bupivacaine. Thus, the overall incidence of females requiring therapeutic agents was quite low.

### Clinical Observations

The results of the clinical observations test did not indicated the presence of acute signs of DA toxicity (vomiting, retching, and seizures) or effects on visual functioning or gross motor coordination. The results did indicate, however, that long-term exposure to DA was related to an increase in observations of subtle arm/hand tremors during the reaching task. Tremors were noted if observed and confirmed on a 2^nd^ trial during the 3 trial sessions. Observers indicated that confirmation of the observation was critical for reliability. The % of total sessions that tremors were observed was calculated over the 2 month baseline period. For the DA exposure period, the % of total sessions that tremors were observed was calculated based on the number of sessions conducted from the beginning of DA exposure for each month up to the time of delivery (see Figure 2). Tremors were rarely observed during the baseline period. The % of total sessions that tremors were observed during the 2-month baseline period was under 20% for all females except one, a female in the control group who exhibited tremors on 26% of the baseline sessions. Tremors increased, however, in a number of females following chronic DA exposure. Tremors were noted on over 20% of the total sessions performed from the 1^st^ day of DA exposure until delivery (or the end of breeding for the 2 females that did not conceive) for 55% (6/11) of females in the 0.15 mg/kg/day DA exposure group, 50% (5/10) of the females in the 0.075 mg/kg/day DA exposure group and only 1 of the 10 (10%) control females (the same female that exhibited tremors for over 20% of the sessions during baseline). The results of a Chi-Square analysis on the distribution of the proportion of total sessions that tremors were noted below and above 20% of the time for all 3 DA exposure groups was close to statistical significance (Χ^2^= 5.17, df=2, p= 0.08). A Chi-Square analysis comparing the controls vs all of the DA exposed females regardless of expousre level indicated that, by the time of delivery, DA exposed females had a higher proportion of total sessions where tremors were noted above 20% of the time (Χ^2^= 5,13, df=1, p= 0.02).

**Figure 2:**
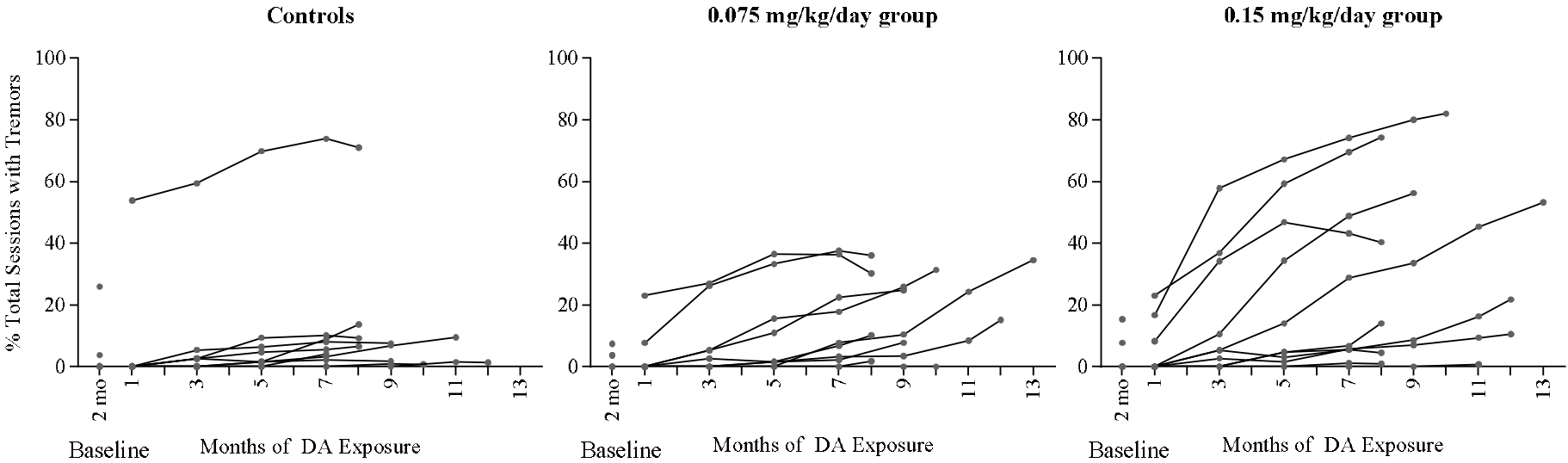
**% of Total Sessions Arm/Hand Tremors Observed on Reaching Task during Baseline and DA Exposure Up To Delivery^1^**

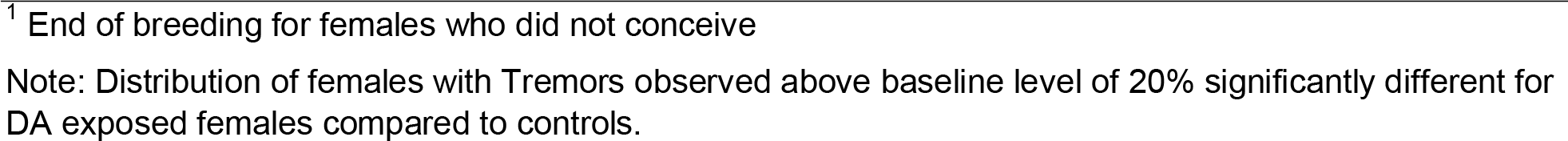

## 3.3 Maternal Reproduction

### Tracking Menstrual Bleeding

All females demonstrated 4 menstrual cycles prior to breeding, 2 cycles during the baseline period and 2 cycles once DA exposure began. There was little change in the length of the menstrual cycle over time. The results of a repeated measures analysis of variance examining changes in the menstrual cycle length for the initial 4 cycles did not indicate a significant main effect of DA exposure on the length of the menstrual cycle (F(2,29) = 1.46, p=0.26), or an interaction of DA exposure and cycle number (1-4) (F(6,87) = 0.84, p=0.54). No main effect of cycle # was observed as well (F(3,87) = 0.09, p=0.97). The average menstrual cycle lengths across the 3 DA exposure groups ranged from 29 to 33 days, normal for fascicularis macaques (see Table 4).

**Table 4:**
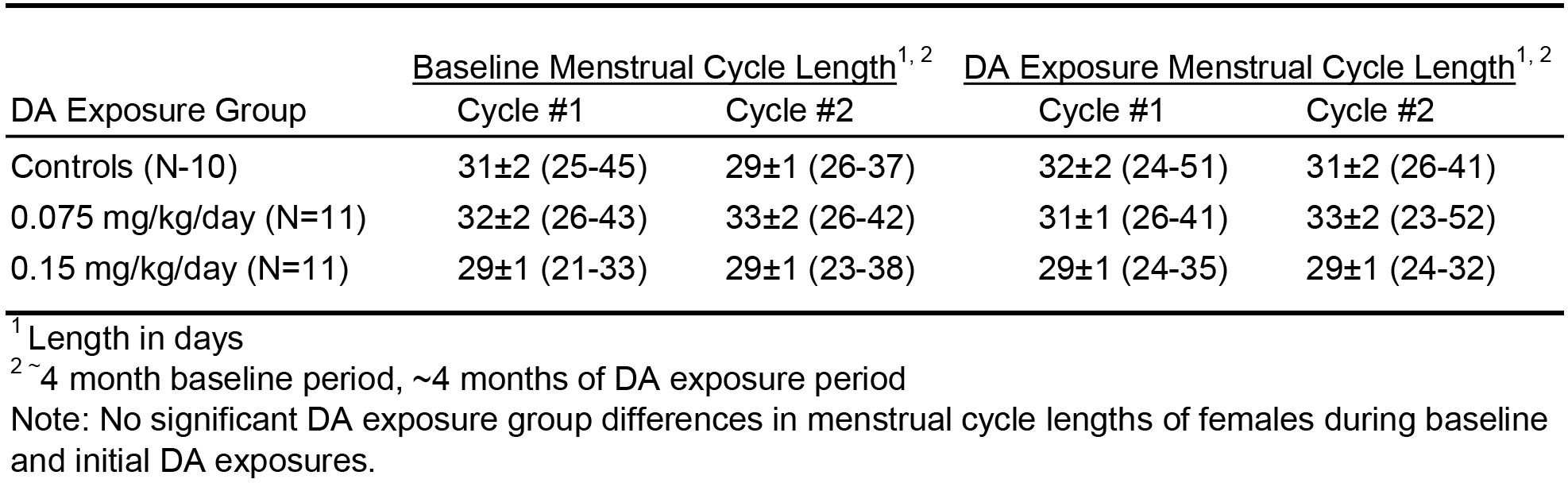
**Mean ± SE (Range) Menstrual Cycle Length Prior to and during DA Exposure**

| DA Exposure Group | Baseline Menstrual Cycle Length<sup>1, 2</sup> |  | DA Exposure Menstrual Cycle Length<sup>1, 2</sup> |  |
| --- | --- | --- | --- | --- |
|  | Cycle #1 | Cycle #2 | Cycle #1 | Cycle #2 |
| Controls (N=10) | 31±2 (25-45) | 29±1 (26-37) | 32±2 (24-51) | 31±2 (26-41) |
| 0.075 mg/kg/day (N=11) | 32±2 (26-43) | 33±2 (26-42) | 31±1 (26-41) | 33±2 (23-52) |
| 0.15 mg/kg/day (N=11) | 29±1 (21-33) | 29±1 (23-38) | 29±1 (24-35) | 29±1 (24-32) |

<sup>1</sup> Length in days
<sup>2</sup> ~4 month baseline period, ~4 months of DA exposure period
Note: No significant DA exposure group differences in menstrual cycle lengths of females during baseline and initial DA exposures.

### Breeding

Conception rates were high (90%-100%) for all DA exposure groups (see Table 5). While the number of breedings required for a conception varied widely (1 to 7), the results of a Chi-Square analysis did not indicate a significant difference in the distribution of the number of breedings to conception due to DA exposure (Χ^2^= 8.78, df=12, p=0.72). The average number of breedings to conception across the exposure groups varied slightly from 2.1 to 2.6.

### Pregnancy and Delivery

Delivery complications occurred frequently in all DA exposure groups, resulting in high cesarean (C-) section delivery rates. Over half of the control and 0.075 mg/kg/day DA females and nearly two-thirds of 0.15 mg/kg/day DA females were delivered via C-section. C-section deliveries were typically related to unproductive labor and/or vaginal bleeding without labor. Decisions regarding C-sections were made by the attending veterinarians from the Primate Center, who were blinded to treatments. The policy of the Primate Center is to err on the side of caution to minimize fetal loss. While the C-section delivery rate was high, so was the livebirth delivery rate, with only 1 pregnancy ending in a nonviable offspring for the entire study (see Table 5). One female in the 0.15 mg/kg/day DA exposure group had a breech delivery. The newborn was delivered via C-section but was not breathing and could not be resuscitated. Nine live-born infants (5 males and 4 females) were delivered in the control group, 9 (5 males and 4 females) in the 0.075 mg/kg/day DA exposure group and 10 (4 males and 6 females) in the 0.15 mg/kg/day DA exposure group.

**Table 5:**
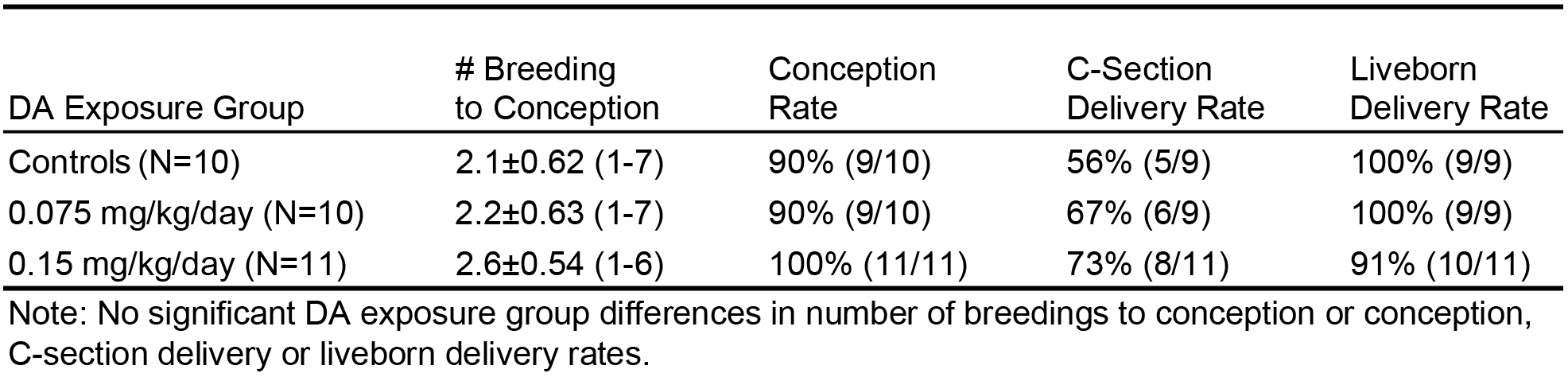
**Mean ± SE (Range) # of Breedings to Conception and Rates for Female Reproductive Outcomes**

Given the high rate of C-sections in the study, a preliminary two-way analysis of variance comparing the length of pregnancy for the females who were C-sectioned to those who delivered vaginally for the DA exposure groups did not indicated a significant main effect of DA exposure group (F(2,23) = 0.69, p=0.51), delivery type (F(1,23) = 0.67, p=0.42) or an interaction of DA exposure group and delivery type (F(2,23) = 1.12, p=0.34). A follow-up one-way ANOVA comparing the length of pregnancies across the 3 DA exposure groups did not indicate a significant difference due to DA exposure (F(2,26) = 0.16, p=0.86). The average pregnancy length across the DA exposure groups varied slightly from 160 to 162 days (see table 6).

**Table 6:** **Mean ± SE (Range) Length of Pregnancy**

| DA Exposure Group | Pregnancy Length |
| --- | --- |
| Controls (N=9) | 162±2.3 (151-172) |
| 0.075 mg/kg/day (N=9) | 162±2.3 (152-170) |
| 0.15 mg/kg/day (N=11) | 160±2.0 (146-170) |

Note: No significant DA exposure group differences in length of pregnancy.

## *3.4* Newborn Procedures

### Newborn Weights and Anthropometries

Birthweights and newborn anthropometric measures are displayed in Table 7. One 0.15 mg/kg/day female infant was vaginally delivered pre-term (<150 days) and had the lowest birthweight at 240 grams. The remaining infants exhibited birthweights and anthropometric measures consistent with normal full-term newborns for this species. The results of two-way ANOVA tests examining the effects of delivery type (C-section and vaginal deliveries) on birthweights and anthropometric measures for the 3 DA exposure groups did not indicate a significant main effect of DA exposure on any of the newborn parameters (p>0.10, all tests), or an interaction of DA exposure and delivery type (p>0.10, all tests). No main effect of delivery type was observed as well (p>0.10, all tests). In addition, results of two-way ANOVA tests examining the effects of sex of the infant on birthweights and anthropometric measures for the 3 DA exposure groups not indicate a significant main effect of DA exposure on any of the newborn parameters (p>0.10, all tests), or an interaction of DA exposure and sex of the infant (p>0.10, all tests). There were, however, significant main effects of sex of the infant on birthweight (F(1,22) = 4.66, p=0.04), head circumference (F(1,22) = 5.90, p=0.02), and head length (F(1,22) = 4.53, p=0.05). Male infants had higher birthweights and larger head circumference and head length measures than females. Three-way ANOVA tests (DA exposure, delivery type and sex of infant) could not be performed due to the low number of infants (N=1) in one of the cells.

**Table 7:** **Mean ± SE (Range) for Infant Birth Characteristics**

**Males**
| DA Exposure Group | Birthweight | Crown-Rump Length | Head Circumference | Head Width | Head Length |
| --- | --- | --- | --- | --- | --- |
| Controls (N=5) | 330±7.2<br>(310-355) | 174.2±3.0<br>(166.5-181.0) | 182.0±1.3<br>(177.0-184.5) | 48.6±0.6<br>(47.0-50.5) | 60.6±0.7<br>(58.0-62.5) |
| 0.075 mg/kg/day (N=5) | 389±26.9<br>(325-480) | 180.3±4.8<br>(168.5-194.0) | 184.2±2.1<br>(178.0-189.5) | 50.0±0.5<br>(48.0-51.0) | 62.0±0.7<br>(60.0-63.5) |
| 0.15 mg/kg/day (N=3) | 332±38.8<br>(255-380) | 177.0±8.5<br>(160.0-186.5) | 183.3±2.4<br>(180.0-188.0) | 48.7±0.3<br>(48.0-49.0) | 60.5±1.0<br>(58.5-61.5) |

**Females**
| DA Exposure Group | Birthweight | Crown-Rump Length | Head Circumference | Head Width | Head Length |
| --- | --- | --- | --- | --- | --- |
| Controls (N=4) | 309±19.0<br>(255-340) | 169.1±4.4<br>(161.5-181.5) | 179.5±3.4<br>(171.5-187.0) | 48.3±1.3<br>(46.0-51.0) | 59.8±1.2<br>(57.0-62.5) |
| 0.075 mg/kg/day (N=4) | 318±20.9<br>(280-365) | 172.8±3.9<br>(164.5-180.5) | 178.4±2.1<br>(172.0-181.0) | 49.1±1.6<br>(47.0-54.0) | 60.3±0.3<br>(59.5-61.0) |
| 0.15 mg/kg/day (N=7) | 304±19.6<br>(240-365) | 173.4±2.8<br>(164.0-183.0) | 177.9±2.0<br>(170.0-182.5) | 47.3±0.6<br>(45.0-49.0) | 58.9±0.7<br>(56.0-60.5) |

Note: Significant differences between males and females for Birthweight ( $p=0.04$ ), Head Circumference ( $p=0.02$ ) and Head Length ( $p=0.05$ ). No significant DA exposure group differences or interaction of DA exposure and sex of the newborn for Infant Birth Characteristics.

### Newborn Health Status (Apgar) and Screening for Congenital Defects

No congenital defects were observed in newborns. The initial Apgar evaluations were performed at the first possible time following delivery, which ranged from 5 to 30 minutes. Follow up evaluations were performed at ~ 1 hour after birth. Apgar scores ranged from 6 to the maximum score of 12 (see Table 8. The results of two-way ANOVAs examining the effects of delivery type (C-section and vaginal deliveries) on the Apgar scores for the 3 DA exposure groups did not indicate a significant main effect of DA exposure for either of the Apgar tests (p>0.10, both tests), or an interaction of DA exposure and delivery type (p>0.10, both tests). There was, however, a significant main effects of delivery type on the initial Apgar scores (F(1,21) = 41.39, p<0.01) and a trend toward an effect on the second Apgar scores (F(1,21) = 3.09, p=0.09). Apgar scores were lower for C-section delivered newborns at both tests. In addition, results of two-way ANOVA tests examining the effects of sex of the infant on Apgar scores did not indicate a significant main effect of DA exposure for either of the Apgar tests (p>0.10, both tests), or an interaction of DA exposure and sex of the infant (p>0.10, both tests). No main effect of sex of the infant was observed as well (p>0.10, both tests).

**Table 8:** **Mean ± SE (Range) Apgar Scores^1^**

**C-Section Delivery**
| DA Exposure Group | Apgar 1 | Apgar 2 |
| --- | --- | --- |
| Controls (N=4) | 7±0.8 (6-12) | 10±0.4 (9-11) |
| 0.075 mg/kg/day (N=3) | 8±0.5 (6-9) | 10±0.4 (9-12) |
| 0.15 mg/kg/day (N=3) | 8±0.6 (6-11) | 10.0±0.6 (7-12) |

**Vaginal Delivery**
| DA Exposure Group | Apgar 1 | Apgar 2 |
| --- | --- | --- |
| Controls (N=5) | 12±0.3 (11-12) | 11±0.4 (10-12) |
| 0.075 mg/kg/day (N=6) | 11±0.3 (11-12) | 11±0.3 (11-12) |
| 0.15 mg/kg/day (N=7) | 11±0.4 (10-11) | 10±1.2 |

<sup>1</sup> Apgar 1 tested ~ 5 to 30 mins after delivery, Apgar 2 tested ~ 1 hour after delivery
<sup>2</sup> N=6
Note: Significant differences between C-Section and Vaginal delivery for Apgar 1 ( $p=0.01$ ), trend for Apgar 2 ( $p=0.09$ ). No significant DA exposure group differences or interaction of DA exposure and delivery type for Apgar scores

### Newborn Plasma DA Concentrations

DA was not detected in the plasma of control females at delivery or their offspring. Plasma DA concentrations at delivery for the adult females in the 0.075 mg/kg/day DA exposure group ranged from 0.24 ng/ml to 4.50 ng/ml. For the adult females in the 0.15 mg/kg/day group, concentrations ranged from 0.74 ng/ml to 5.77 ng/ml. Newborn plasma DA concentrations ranged from 0.24 ng/ml to 0.99 ng/ml for the 0.075 mg/kg/day DA exposure group and 0.31 ng/ml to 3.19 ng/ml for the 0.15 mg/kg/day group (see Table 9). Some of the variability in the plasma DA concentrations is most likely due to the variation in the time between the blood draw and the last maternal dose, which ranged from ~3 to 27 hours for the females and ~4 to 27 hours for the newborns (see Table 9). The results of ANOVA tests indicated both maternal and newborn plasma DA concentrations varied with the timing of the post-delivery blood draw (maternal, F(1,17) = 8.40, p=0.01; newborn, F(1,16) = 8.32, p=0.01). Analysis of co-variance (ANCOVA) tests examining the effects of delivery type (C-section and vaginal deliveries) did not indicate significant effect of this variable on maternal plasma DA concentrations. A follow-up one-way ANCOVA examining the differences in maternal plasma DA concentrations by DA exposure group did not indicate a significant difference between the groups (F(1,16) = 2.48, p=0.14), after taking into account the timing of the blood draw. ANCOVA tests examining the effects of delivery type or sex of the newborn did not indicate significant effects of these variables on newborn plasma DA concentrations. The results of a one-way ANCOVA, however, did indicate that newborn plasma DA concentrations were significantly higher for the 0.15 mg/kg/day DA exposure group compared to the 0.075 mg/kg/day DA exposure group (F(1,15) = 8.49, p=0.01), after taking into account the timing of the blood draw. Finally, an ANCOVA comparing the plasma DA concentrations of the mother versus and newborn indicated that newborn plasma DA concentrations were significantly lower than maternal concentrations for both DA exposure groups (F(1,32) = 7.63, p<0.01),

**Table 9:**
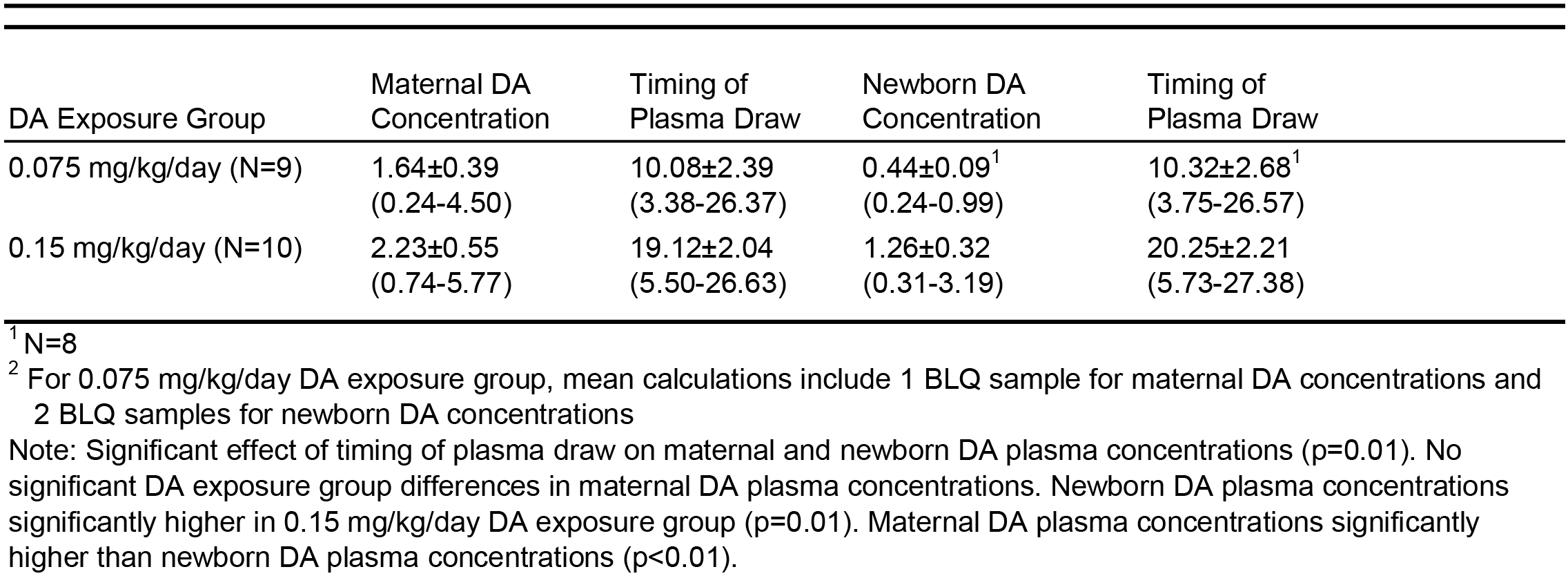
**Mean ± SE (Range) Maternal and Newborn Plasma DA Concentrations (ng/ml) at Delivery and Timing of Plasma Draw (Hours since Last Maternal Dose)**

| DA Exposure Group | Maternal DA Concentration | Timing of Plasma Draw | Newborn DA Concentration | Timing of Plasma Draw |
| --- | --- | --- | --- | --- |
| 0.075 mg/kg/day (N=9) | 1.64±0.39<br>(0.24-4.50) | 10.08±2.39<br>(3.38-26.37) | 0.44±0.09<sup>1</sup><br>(0.24-0.99) | 10.32±2.68<sup>1</sup><br>(3.75-26.57) |
| 0.15 mg/kg/day (N=10) | 2.23±0.55<br>(0.74-5.77) | 19.12±2.04<br>(5.50-26.63) | 1.26±0.32<br>(0.31-3.19) | 20.25±2.21<br>(5.73-27.38) |

<sup>1</sup> N=8
<sup>2</sup> For 0.075 mg/kg/day DA exposure group, mean calculations include 1 BLQ sample for maternal DA concentrations and 2 BLQ samples for newborn DA concentrations
Note: Significant effect of timing of plasma draw on maternal and newborn DA plasma concentrations ( $p=0.01$ ). No significant DA exposure group differences in maternal DA plasma concentrations. Newborn DA plasma concentrations significantly higher in 0.15 mg/kg/day DA exposure group ( $p=0.01$ ). Maternal DA plasma concentrations significantly higher than newborn DA plasma concentrations ( $p < 0.01$ ).

## 4.0 Discussion

This study represents the first scientific effort to chart the reproductive and developmental toxicity of DA in a nonhuman primate model. Although data are available from reports of high-level, acute DA poisoning, very little is known about the biological and behavioral effects of low level, chronic oral DA intake. The study was designed to model environmentally relevant exposure scenarios and focused on the translational value of the preclinical primate model for the protection of human health. Previous reports of DA exposure and effects in humans have relied on consumption measures to estimate exposure since biomarkers of DA were not readily available (Boushey et al., 2016, Grattan et al., 2016, 2018). Thus, little is known regarding the toxicokinetics and resulting body burden of DA following chronic, low-level DA exposure in humans. DA concentrations in the current study were dose-dependent, with significantly higher plasma DA levels in the 0.15 mg/kg/day DA exposure group compared to the 0.075 mg/kg/day group. The results also indicate that, while plasma DA concentrations vary during long-term exposure, there is no evidence of consistent increases in DA levels with prolonged exposure. The DA concentrations in this cohort are consistent with our previous nonhuman primate studies that showed flip-flop kinetics and a relatively long apparent plasma half-life for DA. The data are also consistent with the high inter-individual variability observed in our prior studies (jing et al., 2017). DA was consistently detected in all newborns in the study. The data, however, do not suggest disproportionate accumulation of DA in the newborn after chronic DA exposure. Newborn plasma DA concentrations were consistently lower than those in the dams, suggesting some protective barrier in the placenta that should be identified and characterized. Whether a similar barrier exist in humans is unknown.

Reports of DA toxicity from human poisonings as well as recent epidemiologic studies of coastal communities exposed to DA via dietary intake of shellfish have not investigated reproductive parameters. Our study is the first to pursue these outcomes in a nonhuman primate model. Results from the current study indicate that DA exposure is not associated with female reproductive toxicity; parameters such as the length of the menstrual cycle, the ability to conceive, or the ability to deliver full-term, healthy newborn infants were not compromised in DA exposed females. Studies using rodent models have examined the effects of DA exposure during pregnancy on reproductive success (Levin et al., 2005; Shiotani et al., 2017; Tanemura et al., 2009). The results of these studies also indicate that DA is not a reproductive toxin. In both mouse and rat models of *in utero* exposure, maternal parameters, such as weight gain during pregnancy and litter size, were unaffected. Wildlife studies of feral sea lions, however, have reported increases in abortions and stillbirth in DA exposed animals and reproductive toxicity is a hallmark of a clinical diagnosis of DA toxicosis in these animals (Goldstein et al., 2009).

While the current study did not provide evidence of reproductive toxicity due to chronic, low-level DA exposure, the results did indicate the presence of subtle neurological effects in the form of intentional tremors in the adult females. Intentional tremors were observed when females were extending their arms to actively reach for and pick up a food reward from the hand of one of the research staff. The research staff were reliably trained to detect the presence or absence of these tremors during standardized testing sessions throughout the project. The tremors observed were evident only when females were performing the reaching and pick-up task; no tremors were observed during general observations of females in their cages. One control female exhibited tremors throughout the study; with tremors observed on nearly 70% of the total test sessions at the time of her infant delivery. In DA-exposed subjects, there were significant individual differences in the expression of this treatment effect. For the 0.075 mg/kg/day DA exposure group, 5 of the 10 females rarely exhibited tremors (2% to 15%), while the remaining 5 exhibited tremors on 25% to 35% of test sessions. For the 0.15 mg/kg/day DA exposure group, 5 of the 11 females rarely exhibited tremors (1% to 14%), while 4 females exhibited tremors on 50% to 80% of test sessions.

Wildlife studies of sea lions and studies using rodent and nonhuman primate animal models have reported signs of neurological injury in DA exposed adults. Studies of feral sea lions have reported pacing, ataxia, head weaving and spontaneous recurrent seizures in animals exhibiting DA toxicosis (Cook et al., 2011; Goldstein et al., 2008; Gulland et al., 2002; Scholin et al., 2000). Studies of rodent models have reported persistent neurological injury after DA exposure that is expressed through spontaneous recurrent seizures and epileptic disease (Muha and Ramsdell, 2011; Tiedeken and Ramsdell, 2013). In the rat model, a single IP dose of 4.0 or 7.5 mg/kg DA resulted in generalized tremors, wet-dog shakes, praying posture (anterior part of body raised with front paws clasped) and loss of postural control (Tryphonas et al., 1990c). These high doses ultimately produced status epilepticus and post mortem histopathology showed selective encephalopathy, neuronal degeneration and vacuolation of the neuropil in the limbic (hippocampus, amygdala) and olfactory systems. In the preclinical nonhuman primate model, evidence of DA-induced tremors has been observed in past investigations but only in animals exhibiting DA toxicosis after IV and IP (non-oral) routes of exposure. In macaque monkeys, a single IV dose at 1.0 mg/kg and above (4.0 mg/kg) resulted in tremors and death in 4/7 animals (Scallet et al., 1993). Similarly, Tryphonas and colleagues explored IV and IP DA exposure in macaques and found that single DA doses above 0.5 mg/kg IV and 4.0 mg/kg IP produced tremors, rigidity of movement and loss of balance (Tryphonas et al., 1990a). Lesions consisting of vacuolation of the neuropil, astrocytic swelling, and neuronal shrinkage and hyperchromasia were detected in the area postrema, hypothalamus, hippocampus and inner layers of the retina. In contrast to IV and IP routes of exposure, past studies of oral dosing of up to 10 mg/kg DA have not reported tremors in exposed primate subjects (Truelove et al., 1997; Tryphonas et al., 1990d). While these doses are over 10 times higher than those employed in the present study, the studies did not include standard observations of animals to detect the presence of intentional tremors. These subtle neurological effects, therefore, would have likely gone unnoticed.

Recent results from the epidemiological CoASTAL cohort study in Washington State demonstrate that chronic DA exposure (as measured by the consumption of razor clams) is associated with decrements in memory that are serious enough to affect everyday living skills (Grattan et al., 2018, 2016a). This striking research finding suggests that chronic exposure to low-level DA from harvested shellfish is not without risk to the adult central nervous system. The current study was initiated prior to the publication of the results from the CoASTAL study and focused on evaluating the physical, social and cognitive effects of prenatal DA exposure on the offspring. Assessments of cognition and memory were not included in the study for the DA exposed adult females. Thus, we cannot say at this time whether the tremors represent the sole neurobehavioral effect due to the chronic, low-level DA exposure. We were, however, able to secure supplemental funding based on the observations of tremors in our adult females for further evaluation of a subset of the animals using magnetic resonance based diffusion tensor imaging (DTI) (Petroff et al., under review). DTI can be used to characterize pathologic changes such as ischemia, demyelination, axonal damage, inflammation and edema in the white matter of the brain. A study by Cook et al. (2018) using a post-mortem DTI analysis of the brains of California sea lions diagnosed with DA toxicosis described decreased anisotropy, a measure of axonal and myelin integrity, in the fornix, the white matter tract that is located between the hippocampus and the thalamus. The results of our DTI analysis indicated that increases in intentional tremors were associated with decreased fractional anisotropy in the internal capsule, fornix, pons and corpus callosum. The results could be due to direct damage to the myelin/axonal tracts in these areas or due to the replacement of axonal bundles with other cells (i.e. gliosis) (Alba-Ferrara and de Erausquin, 2013; Budde et al., 2011; Garcia-Lazaro et al., 2016; Smith et al., 2006). Additional post-mortem studies are now planned for the brains of DA-exposed and control females to evaluate histopathological changes in affected brain regions and alterations in gene expression.

The present study is the first to address fetal DA exposure and birth outcomes in a nonhuman primate model. Consistent with past research in other species, DA did not act as a classic teratogen and newborns were free from physical malformations (Levin et al., 2005; Shiotani et al., 2017). There were no exposure-related differences on birth characteristics, but species-typical sex effects on birthweight and head size (males infants were heavier and larger) were observed. An adaptation of the human newborn Apgar assessment was used to evaluate the health of the newborns and no DA related effects were observed. As anticipated, C-sectioned newborns exhibited lower Apgar scores than newborns from natural deliveries (Yeekian et al., 2013). C-sectioned newborns typically exhibited poorer muscle tone and color and were less active than newborns delivered naturally. These effects, however, were short-lived and scores tended to normalize by the 2^nd^ Apgar test approximately 1 hour after delivery. The ultimate goal of the current study, however, is to evaluate the physical and neurobehavioral development of the DA and control offspring. The cohort of infants in this study are currently being evaluated on a targeted series of neurobehavioral and electroencephalogram (EEG) assessments to study the effects of in utero DA exposure on postnatal development. Past studies using rodent models have reported a constellation of effects following prenatal DA exposure that suggest a marked fetal sensitivity to DA, characterized by enduring changes in a number of behavioral domains, particularly memory, activity levels, social behavior and emotional reactivity (for reviews see Costa et al., 2010; Doucette and Tasker, 2016; Grant et al., 2010; Lefebvre and Robertson, 2010). A number of studies have reported dose- and gender-related differences in some of these outcomes (Adams et al., 2009; Baron et al., 2013; Burt et al., 2008; Doucette et al., 2007, 2004; Gill et al., 2012; Levin et al., 2006, 2005; Marriott et al., 2014; Ryan et al., 2011; Shiotani et al., 2017; Zuloaga et al., 2016). Overall, findings with the rodent model suggest that early life DA exposure is linked to significant changes in brain structure and function that do not diminish over time. Future reports from the current investigation will provide the first nonhuman primate data on the risks of in-utero exposure to oral DA within environmentally relevant exposure scenarios and will provide greater scientific resolution on the adequacy of regulatory guidelines to protect the health of high-frequency shellfish consumers and their infants.

## Transparency Document

The transparency document associated with this article can be found in the online version.

## Acknowledgments

This research was supported by grants from the U.S. National Institutes of Health R01 ES023043, P51 OD010425, HD083091 and NCATS Grant TL1 TR000422 (SS). The authors would like to acknowledge the staff and volunteers of the Infant Primate Research Laboratory and the University of Washington National Primate Research Center for their skilled assistance. In addition, we would like to thank Mr. Steven Ellis for his assistance with data processing and analysis.

## References

Adams, A.L., Doucette, T.A., James, R., Ryan, C.L., 2009. Persistent changes in learning and memory in rats following neonatal treatment with domoic acid. Physiol. Behav. 96, 505–12.https://doi.org/10.1016/j.physbeh.2008.11.019

Alba-Ferrara, L.M., de Erausquin, G.A., 2013. What does anisotropy measure? Insights from increased and decreased anisotropy in selective fiber tracts in schizophrenia. Front. Integr. Neurosci. 7, 9. https://doi.org/10.3389/fnint.2013.00009

Apgar,V., 1953. A proposal for a new method of evaluation of the newborn infant, in: Anesthesia and Analgesia. pp. 1056–1059. https://doi.org/10.1213/ANE.0b013e31829bdc5c

Backer, L., Miller, M., 2016. Sentinel Animals in a One Health Approach to Harmful Cyanobacterial and Algal Blooms. Vet. Sci. 3, 8. https://doi.org/10.3390/vetsci3020008

Baron, A.W., Rushton, S.P., Rens, N., Morris, C.M., Blain,P.G., Judge, S.J., 2013. Sex differences in effects of low level domoic acid exposure. Neurotoxicology 34, 1–8. https://doi.org/10.1016/j.neuro.2012.10.010

Bates, S.S., Bird, C.J., de Freitas, A.S.W., Foxall, R., Gilgan, M., Hanic, L., Johnson, G.R., McCulloch, A.W., Odense, P., Pocklington, R., Quilliam, M.A., Sim, G., Smith, J.C., Subba Rao, D.V., Todd, E.C.D., Walter, J.A., Wright, J.L.C., 1989. Pennate Diatom Nitzschia pungens as the Primary Source of Domoic Acid, a Toxin in Shellfish from Eastern Prince Edward Island, Canada. Can. J. Fish Aquat. Sci. 46, 1203–1215.

Berdalet, E., Fleming, L.E., Gowen, R., Davidson, K., Hess, P., Backer, L., Moore, S.K., Hoagland, P., Enevoldsen, H., 2016. Marine harmful algal blooms, human health and wellbeing: Challenges and opportunities in the 21st century. J. Mar. Biol. Assoc. United Kingdom 96, 61–91. https://doi.org/10.1017/S0025315415001733

Bossart, G.D., 2011. Marine mammals as sentinel species for oceans and human health. Vet. Pathol. 48, 676–690. https://doi.org/10.5670/oceanog.2006.77

Boushey, C.J., Delp, E.J., Ahmad, Z., Wang, Y., Roberts, S.M., Grattan, L.M., 2016. Dietary assessment of domoic acid exposure: What can be learned from traditional methods and new applications for a technology assisted device. Harmful Algae 57, 51–55. https://doi.org/10.1016Zj.hal.2016.03.013

Budde, M.D., Janes, L., Gold, E., Turtzo, L.C., Frank, J.A., 2011. The contribution of gliosis to diffusion tensor anisotropy and tractography following traumatic brain injury: Validation in the rat using Fourier analysis of stained tissue sections. Brain 134, 2248–2260. https://doi.org/10.1093/brain/awr161

Burbacher, T., Shen, D., Grant, K., Sheppard, L., Damian, D., Ellis, S., Liberato, N., 1999. Reproductive and offspring developmental effects following maternal inhalation exposure to methanol in nonhuman primates. Res. Rep. Health. Eff. Inst. i-ii, 1-117; discussion 119–133.

Burbacher, T.M., Grant, K.S., Shen, D.D., Sheppard, L., Damian, D., Ellis, S., Liberato, N., 2004. Chronic maternal methanol inhalation in nonhuman primates (Macaca fascicularis): Reproductive performance and birth outcome. Neurotoxicol. Teratol. 26, 639–650. https://doi.org/10.1016/j.ntt.2004.06.001

Burbacher, T.M., Mohamed, M.K., Mottett, N.K., 1987. Methylmercury effects on reproduction and offspring size at birth. Reprod. Toxicol. 1, 267–78.

Burt, M.A., Ryan, C.L., Doucette, T.A., 2008. Altered responses to novelty and drug reinforcement in adult rats treated neonatally with domoic acid. Physiol. Behav. 93, 327–336. https://doi.org/10.1016/j.physbeh.2007.09.003

Buse, E., Habermann, G., Osterburg, I., Korte, R., Weinbauer, G.F., 2003. Reproductive/developmental toxicity and immunotoxicity assessment in the nonhuman primate model. Toxicology 185, 221–227. https://doi.org/10.1016/S0300-483X(02)00614-5

Carter, A.M., 2007. Animal Models of Human Placentation - A Review. Placenta 28. https://doi.org/10.1016/j.placenta.2006.11.002

Centers for Disease Control and Prevention, 2017. Harmful Algal Bloom (HAB)-Associated Illness [WWW Document]. URL https://www.cdc.gov/habs/general.html (accessed 9.28.18).

Cook, P., Reichmuth, C., Gulland, F., 2011. Rapid behavioural diagnosis of domoic acid toxicosis in California sea lions. Biol. Lett. 7, 536–538. https://doi.org/10.1098/rsbl.2011.0127

Cook, P.F., Berns, G.S., Colegrove, K., Johnson, S., Gulland, F., 2018. Postmortem DTI reveals altered hippocampal connectivity in wild sea lions diagnosed with chronic toxicosis from algal exposure. J. Comp. Neurol. 526, 216–228. https://doi.org/10.1002/cne.24317

Costa,L.G., Giordano, G., Faustman, E.M., 2010. Domoic acid as a developmental neurotoxin. Neurotoxicology 31, 409–423. https://doi.org/10.1016/j.neuro.2010.05.003

Dakshinamurti, K., Sharma, S.K., Sundaram, M., Watanabe, T., 1993. Hippocampal changes in developing postnatal mice following intrauterine exposure to domoic acid. J. Neurosci. 13, 4486–4495.

Doucette, T.A., Bernard, P.B., Husum, H., Perry, M.A., Ryan, C.L., Tasker, R.A., 2004. Low doses of domoic acid during postnatal development produce permanent changes in rat behaviour and hippocampal morphology. Neurotox. Res. 6, 555–563. https://doi.org/10.1007/BF03033451

Doucette, T.A., Ryan, C.L., Tasker, R.A., 2007. Gender-based changes in cognition and emotionality in a new rat model of epilepsy. Amino Acids 32, 317–322. https://doi.org/10.1007/s00726-006-0418-7

Doucette, T.A., Tasker, R.A., 2016. Perinatal domoic acid as a neuroteratogen, in: Current Topics in Behavioral Neurosciences. Springer, Cham, pp. 87–110. https://doi.org/10.1007/7854_2015_417

FAO (Food and Agriculture Organization). 2004. Report of the Joint FAO/IOC/WHO ad hoc Expert Consultation on Biotoxins in Bivalve Mollusks. Oslo, Norway: Sept 26–30.

Ferriss, B.E., Marcinek, D.J., Ayres, D., Borchert, J., Lefebvre, K.A., 2017. Acute and chronic dietary exposure to domoic acid in recreational harvesters: A survey of shellfish consumption behavior. Environ. Int. 101, 70–79.https://doi.org/10.1016yj.envint2017.01.006

Fialkowski, M.K., McCrory, M.A., Roberts, S.M., Tracy, J.K., Grattan, L.M., Boushey, C.J., 2010. Evaluation of dietary assessment tools used to assess the diet of adults participating in the Communities Advancing the Studies of Tribal Nations Across the Lifespan cohort. J. Am. Diet. Assoc. 110, 65–73. https://doi.org/10.1016/jjada.2009.10.012

Garcia-Lazaro, H.G., Becerra-Laparra, I., Cortez-Conradis, D., Roldan-Valadez, E., 2016. Global fractional anisotropy and mean diffusivity together with segmented brain volumes assemble a predictive discriminant model for young and elderly healthy brains: A pilot study at 3T. Funct. Neurol. 31, 39–46. https://doi.org/10.11138/FNeur/2016.31.1039

Garrison, D.L., Conrad, S.M., Eilers, P.P., Waldron, E.M., 1992. Confirmation of Domoic Acid Production By Pseudonitzschia-Australis (Bacillariophyceae) Cultures. J. Phycol. 28, 604–607. https://doi.org/10.1111/j.0022-3646.1992.00604.x

Gill, D.A., Perry, M.A.,McGuire, E.P., Perez-Gomez, A., Tasker, R.A., 2012. Low-dose neonatal domoic acid causes persistent changes in behavioural and molecular indicators of stress response in rats. Behav. Brain Res. 230, 409–417. https://doi.org/10.1016/j.bbr.2012.02.036

Goldstein, T., Mazet, J.K., Zabka, T.S., Langlois, G., Colegrove, K.M., Silver, M., Bargu, S., Van Dolah, F.M., Conrad, P. a, Barakos, J., Williams, D.C., Dennison, S., Haulena, M., 2008. Novel symptomatology and changing epidemiology of domoic acid toxicosis in California sea lions (Zalophus californianus): an increasing risk to marine mammal health. Proc. R. Soc. 275, 267–276. https://doi.org/10.1098/rspb.2007.1221

Goldstein, T., Zabka, T.S., Delong, R., Wheeler, E.A., Ylitalo, G., Bargu, S., Silver, M., Leighfield, T., Van Dolah, F.M., Langlois, G., Sidor, I., Dunn, J.L., Gulland, F.M., 2009. The Role of Domoic Acid in Abortion and Premature Parturition of California Sea Lions (Zalophus Californianus) on San Miguel Island, California. J. Wildl. Dis. 45, 91–108.

Grant, K.S., Burbacher, T.M., Faustman, E.M., Grattan, L.M., 2010. Domoic acid: Neurobehavioral consequences of exposure to a prevalent marine biotoxin. Neurotoxicol. Teratol. 32, 132–141. https://doi.org/10.1016/j.ntt.2009.09.005

Grattan, L.M., Boushey, C.J., Liang, Y., Lefebvre, K.A., Castellon, L.J., Roberts, K.A., Toben, A.C., Morris, J.G.J., 2018. Repeated Dietary Exposure to Low Levels of Domoic Acid and Problems with Everyday Memory: Research to Public Health Outreach. Toxins (Basel). 10, 103. https://doi.org/10.3390/toxins10030103

Grattan, L.M., Boushey, C.J., Tracy, K., Trainer, V.L., Roberts, S.M., Schluterman, N., Morris, J.G.J., 2016a. The association between razor clam consumption and memory in the CoASTAL cohort. Harmful Algae 57, 20–25. https://doi.org/10.1016yj.hal.2016.03.011

Grattan, L.M., Holobaugh, S., Morris, J.G.J., 2016b. Harmful algal blooms and public health. Harmful Algae. https://doi.org/10.1016/j.hal.2016.05.003

Gulland, F.M.D., Haulena, M., Fauquier, D., Langlois, G., Lander, M.E., Zabka, T.S., Duerr, R., 2002. Domoic acid toxicity in Californian sea lions (Zalophus californianus): clinical signs. Vet. Rec. 150, 475–480. https://doi.org/doi:10.1136/vr.150.15.475

Jeffery, B., Barlow, T., Moizer, K., Paul, S., Boyle, C., 2004. Amnesic shellfish poison. Food Chem. Toxicol. 42, 545–557. https://doi.org/10.1016/j.fct.2003.11.010

King, B.F., 1993. Development and structure of the placenta and fetal membranes of nonhuman primates. J. Exp. Zool. 266, 528–540. https://doi.org/10.1002/jez.1402660605

Kirkley, K.S., Madl, J.E., Duncan, C., Gulland, F.M., Tjalkens, R.B., 2014. Domoic acid-induced seizures in California sea lions (Zalophus californianus) are associated with neuroinflammatory brain injury. Aquat. Toxicol. 156, 259–268. https://doi.org/10.1016/j.aquatox.2014.09.003

Lefebvre, K.A., Bargu, S., Kieckhefer, T., Silver, M.W., 2002. From sandabs to blue whales: the pervasiveness of domoic acid. Toxicon 40, 971–977.

Lefebvre, K.A., Kendrick, P.S., Ladiges, W., Hiolski, E.M., Ferriss, B.E., Smith, D.R., Marcinek, D.J., 2017. Chronic low-level exposure to the common seafood toxin domoic acid causes cognitive deficits in mice. Harmful Algae 64, 20–29. https://doi.org/10.1016/j.hal.2017.03.003

Lefebvre, K.A., Quakenbush, L., Frame, E., Huntington, K.B., Sheffield, G., Stimmelmayr, R., Bryan, A., Kendrick, P., Ziel, H., Goldstein, T., Snyder, J.A., Gelatt, T., Gulland, F.M.D., Dickerson, B., Gill, V., 2016. Prevalence of algal toxins in Alaskan marine mammals foraging in a changing arctic and subarctic environment. Harmful Algae 55, 13–24. https://doi.org/10.1016/j.hal.2016.01.007

Lefebvre, K.A., Robertson, A., 2010. Domoic acid and human exposure risks: A review. Toxicon 56, 218–230. https://doi.org/10.1016/j.toxicon.2009.05.034

Levin, E.D., Pang, W.G., Harrison, J., Williams, P., Petro, A., Ramsdell, J.S., 2006. Persistent neurobehavioral effects of early postnatal domoic acid exposure in rats. https://doi.org/10.1016/j.ntt.2006.08.005

Levin, E.D., Pizarro, K., Pang, W.G., Harrison, J., Ramsdell, J.S., 2005. Persisting behavioral consequences of prenatal domoic acid exposure in rats. Neurotoxicol. Teratol. 27, 719–725. https://doi.org/10.1016/j.ntt.2005.06.017

Mariën, K., 1996. Establishing tolerable dungeness crab (Cancer magister) and razor clam (Siliqua patula) domoic acid contaminant levels. Environ. Health Perspect. 104, 1230–6. https://doi.org/10.1289/ehp.961041230

Marriott, A.L., Tasker, A., Ryan, C.L., Doucette, T.A., 2014. Neonatal domoic acid abolishes latent inhibition in male but not female rats and has differential interactions with social isolation. Neurosci. Lett. 578, 22–26. https://doi.org/10.1016/j.neulet.2014.06.025

McCabe, R.M., Hickey, B.M., Kudela, R.M., Lefebvre, K.A., Adams, N.G., Bill, B.D., Gulland, F.M.D.D. Thomson, R.E., Cochlan, W.P., Trainer, V.L., 2016. An unprecedented coastwide toxic algal bloom linked to anomalous ocean conditions. Geophys. Res. Lett. 43, 10,366–10,376. https://doi.org/10.1002/2016GL070023

McKibben, S.M., Peterson, W., Wood, A.M., Trainer, V.L., Hunter, M., White, A.E., 2017. Climatic regulation of the neurotoxin domoic acid. PNAS 114, 239–244. https://doi.org/10.1073/pnas.1606798114

Montie, E.W., Wheeler, E., Pussini, N., Battey, T.W.K., Barakos, J., Dennison, S., Colegrove, K., Gulland, F., 2010. Magnetic resonance imaging quality and volumes of brain structures from live and postmortem imaging of California sea lions with clinical signs of domoic acid toxicosis. Dis. Aquat. Organ. 91, 243–256. https://doi.org/10.3354/dao02259

Muha, N., Ramsdell, J.S., 2011. Domoic acid induced seizures progress to a chronic state of epilepsy in rats. Toxicon 57, 168–171. https://doi.org/10.1016/j.toxicon.2010.07.018

Perl, T.M., Bedard, L., Kosatsky, T., Hockin, J.C., Todd, E.C., McNutt, L.A., Remis, R.S., 1990a. Amnesic shellfish poisoning: a new clinical syndrome due to domoic acid. Canada Dis. Wkly. Rep. 16 Suppl 1, 7–8.

Perl, T.M., Bedard, L., Kosatsky, T., Hockin, J.C., Todd, E.C.D., Remis, R.S., Bédard, L., 1990b. An Outbreak of Toxic Encephalopathy Caused by Eating Mussels Contaminated with Domoic Acid. N. Engl. J. Med. 322, 1775–1780. https://doi.org/10.1056/NEJM199006213222504

Petroff, R., Richards, T., Crouthamel, B., Stanley, C., McKain, N., Grant, K., Burbacher, T., under review.Chronic, Low-Level Oral Exposure to Marine Toxin, Domoic Acid, Alters Whole Brain Morphometry in Nonhuman Primates. Neurotoxicology.

Ramsdell, J.S., Gulland, F.M., 2014. Domoic acid epileptic disease. Mar. Drugs 12, 1185–1207. https://doi.org/10.3390/md12031185

Ryan, C.L., Robbins, M. a, Smith, M., Gallant, I.C.,Adams-Marriott, A.L., Doucette, T.A., 2011. Altered social interaction in adult rats following neonatal treatment with domoic acid. Physiol. Behav. 102, 291–295. https://doi.org/10.1016/j.physbeh.2010.11.020

Scallet, A., Kowalke, P.K., Rountree, R.L., Thorn, B.T., Binienda, Z.K., 2004. Electroencephalographic, behavioral, and c-fos responses to acute domoic acid exposure. Neurotoxicol. Teratol. 26, 331–342. https://doi.org/10.1016/j.ntt.2003.10.004

Scallet, A.C., Binienda, Z.K., Caputo, F.A.A., Hall, S., Paule, M.G., Rountree, R.L., Schmued, L.C., Sobotka, T.J., Slikker, W., 1993. Domoic acid-treated cynomolgus monkeys (M. fascicularis): Effects of Dose on Hippocampal Neuronal and Terminal Degeneration. Brain Res. 627, 307–313. https://doi.org/10.1016/0006-8993(93)90335-K

Scholin, C.A., Gulland, F.M.D., Doucette, G.J., Benson, S., Busman, M., Chavez, F.P., Cordaro, J., DeLong, R., De Vogelaere, A., Harvey, J., Haulena, M., Lefebvre, K.A., Lipscomb, T., Loscutoff, S., Lowenstine, L.J., Marin, R., Miller, P.E., McLellan, W.A., Moeller, P.D.R., Powell, C.L., Rowles, T.T.K., Silvagni, P., Silver, M.W., Spraker, T., Trainer, V.L., Van Dolah, F.M., 2000. Mortality of sea lions along the central California coast linked to a toxic diatom bloom. Nature 403, 80–84. https://doi.org/10.1038/47481

Shiotani,M., Cole, T.B., Hong, S., Park, J.J.Y., Griffith, W.C., Burbacher, T.M., Workman, T., Costa, L.G., Faustman, E.M., 2017. Neurobehavioral assessment of mice following repeated oral exposures to domoic acid during prenatal development. Neurotoxicol. Teratol. 64, 8–19. https://doi.org/10.1016/J.NTT.2017.09.002

Shum, S., Kirkwood, J.S., Jing, J., Petroff, R., Crouthamel, B., Grant, K.S., Burbacher, T.M., Nelson, W.L., Isoherranen, N., 2018. Validated HPLC-MS/MS Method To Quantify Low Levels of Domoic Acid in Plasma and Urine after Subacute Exposure. ACS Omega 3, 12079–12088. https://doi.org/10.1021/acsomega.8b02115

Smith, S.M., Jenkinson, M., Johansen-Berg, H., Rueckert, D., Nichols, T.E., Mackay, C.E., Watkins, K.E., Ciccarelli, O., Cader, M.Z., Matthews, P.M., Behrens, T.E.J., 2006. Tract based spatial statistics: voxelwise analysis of multi-subjects diffusion data. Neuroimage 31, 1487–1505. https://doi.org/10.1016/j.neuroimage.2006.02.024

Stommel, E.W., Watters, M.R., 2004. Marine neurotoxins: Ingestible toxins. Curr. Treat. Options Neurol. https://doi.org/10.1007/s11940-004-0020-9

Tanemura, K., Igarashi, K., Matsugami, T.-R., Aisaki, K., Kitajima, S., Kanno, J., 2009. Intrauterine environment-genome interaction and Children’s development (2): Brain structure impairment and behavioral disturbance induced in male mice offspring by a single intraperitoneal administration of domoic acid (DA) to their dams. J. Toxicol. Sci. 34, SP279–SP286. https://doi.org/10.2131/jts.34.SP279

Teitelbaum, J., Zatorre, R.J., Carpenter, S., Gendron, D., Cashman, N.R., 1990a. Neurological Sequelae of Domoic Acid Intoxication. Symp. Domoic Acid Toxic. 16, 9–12. https://doi.org/10.2174/13816128236661701241

Teitelbaum, J., Zatorre, R.J., Carpenter, S., Gendron, D., Evans, A., Gjedde, A., Cashman, N., 1990b. Neurologic Sequelae of Domoic Acid Intoxication due to the Ingestion of Contaminated Mussels. N. Engl. J. Med. 322, 1781–1787.

Thompson, B.L., Levitt, P., Stanwood, G.D., 2009. Prenatal exposure to drugs: effects on brain development and Implications for Policy and Education. Nat. Rev. Neurosci. 10, 303–312. https://doi.org/10.1038/nrn2598.Prenatal

Tiedeken, J.A., Ramsdell, J.S., 2013. Persistent neurological damage associated with spontaneous recurrent seizures and atypical aggressive behavior of domoic acid epileptic disease. Toxicol. Sci. 133, 133–143. https://doi.org/10.1093/toxsci/kft037

Toyofuku, H., 2006. Joint FAO/WHO/IOC activities to provide scientific advice on marine biotoxins (research report). Mar. Pollut. Bull. 52, 1735–1745. https://doi.org/10.1016/j.marpolbul.2006.07.007

Truelove, J., Mueller, R., Pulido, O., Martin, L., Fernie, S., Iverson, F., 1997. 30-day oral toxicity study of domoic acid in cynomolgus monkeys: Lack of overt toxicity at doses approaching the acute toxic dose. Nat. Toxins 5, 111–114. https://doi.org/10.1002/nt.5

Tryphonas, L., Truelove, J., Iverson, F., 1990a. Acute parenteral neurotoxicity of domoic acid in cynomolgus monkeys (M. fascicularis). Toxicol. Pathol. 18, 297–303. https://doi.org/10.1177/019262339001800208

Tryphonas, L., Truelove, J., Iverson, F., Todd, E.C.D., Nera, E., 1990b. Neuropathology of Experimental Domoic Acid Poisoning in Nonhuman Primates and Rats. Symp. Domoic Acid Toxic. 78–81.

Tryphonas, L., Truelove, J., Nera, E., Iverson, F., 1990c. Acute Neurotoxicity of Domoic Acid in the Rat. Toxicol. Pathol. 18, 1–9. https://doi.org/10.1177/019262339001800101

Tryphonas, L., Truelove, J., Todd, E.C.D., Nera, E., Iverson, F., 1990d. Experimental oral toxicity of domoic acid in cynomolgus monkeys (Macaca fascicularis) and rats. Food Chem. Toxicol. 28, 707–715. https://doi.org/10.1016/0278-6915(90)90147-F

US Department of Commerce, National Oceanic and Atmospheric Administration, n.d. What percentage of the American population lives near the coast?

Washington State Department of Health, n.d. Amnesic Shellfish Poisoning (ASP) from Domoic Acid [WWW Document]. URL https://www.doh.wa.gov/CommunityandEnvironment/Shellfish/RecreationalShellfish/Illnesses/Biotoxins/AmnesicShellfishPoisoning (accessed 9.28.18).

Wekell, J.C., Jurst, J., Lefebvre, K. a, 2004. The origin of the regulatory limits for PSP and ASP toxins in shellfish. J. Shellfish Res. 23, 927–930.

Wells, M.L., Trainer, V.L., Smayda, T.J., Karlson, B.S.O., Trick, C.G., Kudela, R.M., Ishikawa, A., Bernard, S., Wulff, A., Anderson, D.M., Cochlan, W.P., 2015. Harmful algal blooms and climate change: Learning from the past and present to forecast the future. Harmful Algae 49, 68–93. https://doi.org/10.1016/j.hal.2015.07.009

Xi, D., Peng, Y.G., Ramsdell, J.S., 1997. Domoic acid is a potent neurotoxin to neonatal rats. Nat. Toxins 5, 74–79. https://doi.org/10.1002/(SICI)(1997)5:2<74::AID-NT4>3.0.CO;2-I

Yeekian, C., Jesadapornchai, S., Urairong, K., Santibenjakul, S., Suksong, W., Nuchprayoon, C., 2013. Comparison of maternal factors and neonatal outcomes between elective cesarean section and spontaneous vaginal delivery. J. Med. Assoc. Thai. 96, 389–94.

Zhu, Z., Qu, P., Fu, F., Tennenbaum, N., Tatters, A.O., Hutchins, D.A., 2017. Understanding the blob bloom: Warming increases toxicity and abundance of the harmful bloom diatom Pseudo-nitzschia in California coastal waters. Harmful Algae 67, 36–43. https://doi.org/10.1016/j.hal.2017.06.004

Zuloaga, D.G.G., Lahvis, G.P.P., Mills, B., Pearce, H.L.L., Turner, J., Raber, J., 2016. Fetal domoic acid exposure affects lateral amygdala neurons, diminishes social investigation and alters sensory-motor gating. Neurotoxicology 53, 132–140. https://doi.org/10.1016/j.neuro.2016.01.007

